# Task-specificity of local load feedback depends on intersegmental information in an insect walking system

**DOI:** 10.1101/2025.04.29.651003

**Authors:** Angelina Ruthe, Alkan Özyer, Turgay Akay, Ansgar Büschges

**Author notes:** These authors contributed equally; shared first authorship (Angelina Ruthe, Alkan Özyer).

## Abstract

Load proprioceptive feedback measured by campaniform sensilla (CS) substantially contributes to the generation of leg stepping during walking. In stick insects, the effect of load feedback on the antagonistic protractor and retractor motor neurons (MNs) reverses depending on the walking direction to support the functional stance MN pool in both stepping directions. However, this was shown in highly reduced preparations in which the stimulated leg did not perform actual stepping movements, but the effect was inferred from MN activity profiles. It is still largely unknown whether the load feedback similarly shows a direction dependent reversal in its effects on protractor and retractor activity during actual stepping of the stimulated leg. Here, we implemented load stimulation in a single-middle leg preparation and recorded protractor and retractor MN activity extracellularly during fictive forward and backward stepping. We show that CS stimulation during the stance phase prolonged stance (retractor) activity for the duration of the stimulation and induced stance (retractor) activity when applied during the swing phase in forward stepping. During backward stepping, when protractor and retractor switch phases, the same stimulation again prolonged stance (protractor) activity for the duration of the stimulation. However, it rarely induced stance (protractor) activity when applied during the swing phase. We provide evidence that this is due to the absence of signals about the stepping direction of the neighboring legs, thus highlighting the importance of intersegmental signals for task-specific processing of load proprioceptive feedback.

**New and Noteworthy:** Proprioceptive feedback needs to be processed in a task-dependent manner to serve its function, even when the leg kinematics change to generate different behaviors. We show for forward and backward stepping that load signals initiate and maintain the activity of the functional stance phase motor neuron pool and that intersegmental signals about stepping of the neighboring legs are relevant for the task-specific processing of load feedback.

## Introduction

Proprioceptive feedback plays a pivotal role in generating a functional walking motor output in legged animals, vertebrates and invertebrates alike (Orlovsky et al. 1999; Akay et al. 2014; Büschges and Ache 2025; Goulding et al. 2025). In insects, particularly information about forces generated by the leg muscles and load signals from the legs are highly relevant for the generation of a proper motor output during a leg’s stance phase when an appendage contributes individually to moving the body in the desired direction (Zill and Moran 1981; Schmitz 1993; Noah et al. 2004; Zill et al. 2004; Gruhn et al. 2016). Such forces and load are perceived by campaniform sensilla (CS), mechanoreceptors on the leg that are often arranged in groups near the leg joints and respond to strain in the cuticle (Pringle 1938; Chapman et al. 1973; Spinola and Chapman 1975; Zill and Moran 1981; Zill et al. 2012). The neuronal pathways mediating load feedback include direct and polysynaptic pathways between CS and motor neurons (MNs) modulating the activity of MNs in a reflex-like fashion (Zill et al. 1981; Parker and Newland 1995; Schmitz and Stein 2000; Höltje and Hustert 2003; Akay et al. 2007; Haberkorn et al. 2019; Gebehart et al. 2021), and pathways targeting the joint-specific premotor central pattern generating circuits (CPGs; Büschges et al. 1995; Akay et al. 2001, 2007). Together, these influences are known to determine and control the motor activity of the leg during stepping (reviewed in: Zill et al. 2004).

At present, the available knowledge about the influence of proprioceptive feedback signals from a leg on its motor activity can explain the generation of a default stepping mode, such as straight-forward stepping (reviewed in: Büschges and Ache 2025). In this context, feedback signals about the increase and endurance of the load from the leg initiate and maintain the activity of the MNs innervating the stance muscles in the middle leg, for example, the activity of the retractor coxae (retractor) and depressor trochanteris MNs (Akay et al. 2001, 2007; Borgmann et al. 2011). Additionally, evidence is accumulating that proprioceptive feedback needs to be modified when it comes to task-specific changes in leg kinematics, for example, to generate forward or backward stepping or turning (Dürr and Ebeling 2005; Akay et al. 2007; Mu and Ritzmann 2008a; Hellekes et al. 2012; Gruhn et al. 2016). When walking a curve, leg kinematics inside and outside of the curve radius are modified in a way that the outside legs push the animal into the curve, while inside legs modify their leg stepping kinematics to pull the animal towards the curvature (Gruhn et al. 2009; Yang et al. 2024). For forward and backward walking, each leg changes its kinematics to generate stance phases in opposite directions (stick insect: Graham and Epstein 1985; ant: Pfeffer et al. 2016; fruit fly: Bidaye et al. 2014; Feng et al. 2020). For the middle leg of the stick insect, this change is brought about by reversing the activity of those two leg muscles, which move the thoraco-coxal (ThC) joint: the retractor generates stance for forward stepping, and it is the antagonist, the protractor coxae (protractor), who serve stance generation in backward stepping (Graham and Epstein 1985; Rosenbaum et al. 2010, 2015). Previous studies have shown that task-specific influences of proprioceptive feedback contribute to changes in leg kinematics (Hellekes et al. 2012; Gruhn et al. 2016). For example, while feedback signals from load sensors on the legs, the CS, support retractor MN activity in forward walking, they exert the same supporting influence onto protractor MN for backward stepping (Akay et al. 2007). This change of influence is mediated by reversing the influence of load feedback signals on the CPG governing the ThC joint MNs, allowing the conclusion that load signals assist the generation of leg stance for either walking direction (Akay et al. 2007).

One confounding factor of the previous studies is that proprioceptive feedback influences were most often studied in highly reduced preparations. In the case of load feedback, for example, preparations were used, in which advantageously individual CS fields could be stimulated. However, in these experiments, only parts of the leg housing these sensory organs were left intact with no actual movement being generated (Akay et al. 2007; Gruhn et al. 2016). Consequently, relating the influence of sensory stimulation to specific phases of the leg stepping cycle, i.e., stance or swing, was limited to the activity of MN pools related to stance or swing. However, we reasoned that in the presence of actual movement inducing feedback from multiple sensory sources might alter the effect of isolated CS feedback in reduced preparations. Here, we chose to study the influence of load feedback in a stick insect preparation, which was able to generate active stepping movements of the stimulated leg, the so-called *single stepping leg preparation* of the stick insect (Karg et al. 1991; Büschges and Gruhn 2020). This preparation was combined with stimulating trochanteral/femoral campaniform sensilla (tr/fCS) (Hofmann and Bässler 1982, 1986) to analyze the contribution of load feedback signals to the motor activity of retractor and protractor MNs throughout the stepping cycle. Given that stimulating load sensors during semi-intact leg stepping represents a challenging task, we first set up an experimental approach, with which we were able to apply additional load to the stepping leg in a defined way, while simultaneously recording the activity of the coxal MN pools for both stepping directions.

We show here that in forward stepping, load feedback initiates and supports retractor MN activity and thus mediates a stance motor output during both stance and swing phases of the leg stepping cycle. Unloading feedback signals had the opposite effect, supporting the initiation of leg swing motor activity. On the contrary, only a supporting influence of load feedback signals was detected during stance in backward stepping, i.e., prolonging protractor MN activity. Load feedback during swing only rarely initiated protractor and thus stance activity. Interestingly, the influence of load feedback signals during swing in backward stepping appeared to depend on the presence of other stepping legs, and the initiation of stance was only as frequent as reported by Akay et al. (2007) when five other legs were also stepping. Our findings give rise to the conclusion that intersegmental information transfer is relevant for promoting task-specific processing of local feedback signals, which in turn contributes to a precise motor output of the multi-segmented insect leg.

## Materials and Methods

All experiments were performed on adult female stick insects (*Medauroidea extradentata)*. The insects were taken from parthenogenetic breeding colonies at the Biocenter, University of Cologne. The insects were kept at approximately 24 – 26°C, 60 – 65% humidity, 12/12h light/dark cycles, and were fed with blackberry leaves *ad libitum*. The experiments were performed in daylight and at room temperature (20 - 22°C).

### Preparation

#### 1. Single middle leg stepping preparation

The preparation used was a single-leg preparation (Fig. 1A; Fischer et al. 2001; Gabriel and Büschges 2007; Rosenbaum et al. 2015), allowing the leg to perform stepping movements on a treadmill oriented perpendicular to the insect at the level of the middle leg. For this, all legs were removed except for the right middle leg. In this remaining leg, the ThC joint was embedded in dental cement (Protemp II, 3M ESPE) to prevent it from moving. For the fixation, the leg was oriented perpendicular to the body axis (Rosenbaum et al. 2015), and special care was taken not to cover the tr/fCS with the dental cement. This fixation of the ThC joint allows the leg to be moved in the vertical plane. The insect was subsequently glued to the edge of a foam platform with its dorsal side facing upwards. The thorax was opened along the midline of the tergum and fixed with insect pins to gain access to the ventral nerve cord. The gut was moved out of the body cavity and fixed next to the insect. The body cavity was subsequently filled with saline (Weidler and Diecke 1969), and trachea, fat, and connective tissue covering the target nerves were removed. Under the preceding conditions, the middle leg was able to perform fictive stepping movements on a treadmill. The treadmill used in the experiments is custom-made and consists of two Styrofoam drums with a diameter of around 25 mm and a center distance of around 50 mm, connected via a crepe paper belt. The two Styrofoam drums were connected to a DC motor from which the velocity with which the paper belt moved was measured (amplified 50-fold). The height and the position of the treadmill were adjusted to the length of the leg, in particular the length of the tibia, to ensure proper orientation of the leg. A paintbrush was used to stimulate the animal. Forward stepping was induced by gently touching the abdomen and backward stepping was induced by touching the head or antenna regions (Graham and Epstein 1985).

**Figure 1:**
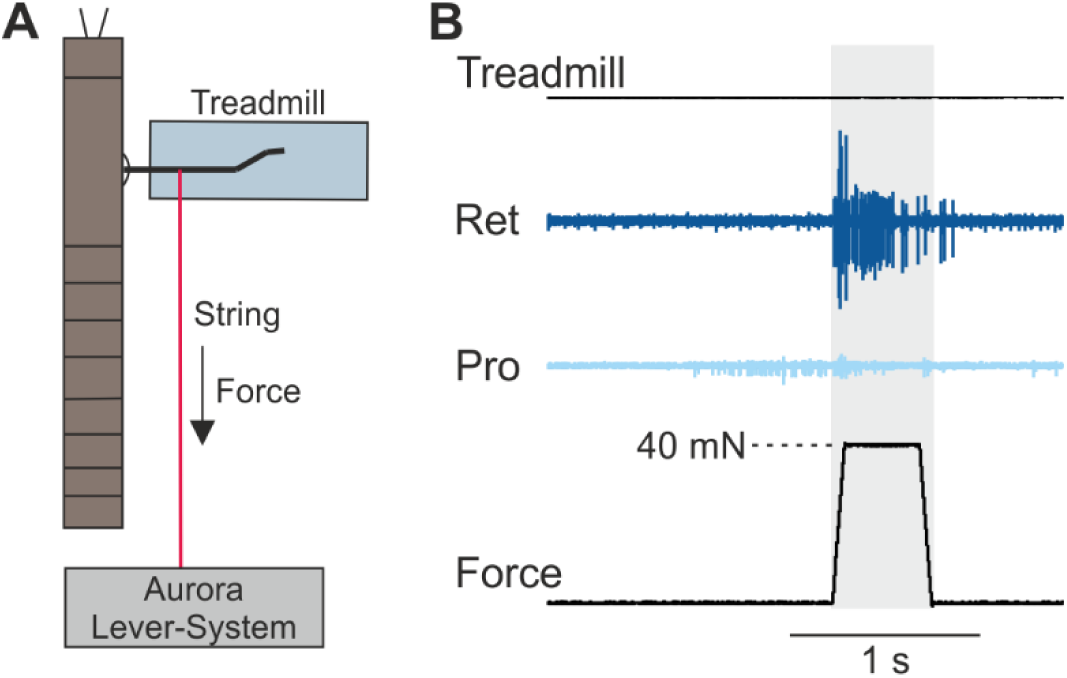
Experimental setup with stimulation of load sensors in the single-middle leg preparation. **A:** Schematic of the single middle leg preparation with stimulation of load sensors. A string is connected to the femur and to a Dual-Mode-Lever-System. 40 mN force is applied to the leg by pulling it in posterior direction. **B:** Extracellular recordings of protractor and retractor MNs during stimulation of load sensors. The upper trace shows the treadmill movement, the bottom trace shows the forces applied to the leg. Ramp-and-hold stimuli of 40 mN with a holding duration of 400 ms were used. MNs: motor neurons, Pro: protractor coxae, Ret: retractor coxae.

#### 2. Preparation with five or less legs stepping on a slippery surface

In this approach, adapted from (Graham and Epstein 1985; Gruhn et al. 2011), five or less legs were able to move freely on an acrylic slippery surface while one was placed on a treadmill. The treadmill was localized at the level of the right middle leg, and up to three individual acrylic slippery surfaces were positioned at the level of the intact moving legs, i.e., one on the contralateral side for the three legs to move, and two smaller ones, one each for the ipsilateral front leg and hind leg. Only in experiments with five legs stepping, a slippery surface was used for the stimulated middle leg as well due to the limited space. Additionally, in experiments with two stepping legs, either two treadmills or one treadmill and one slippery surface were used. These data were pooled for the analysis. The insect was mounted on a balsa rod and prepared for electrophysiological recordings as described in 1. To elicit forward walking, the insect was stimulated with a paintbrush at the abdomen, while for backward stepping, the insect was stimulated at the head region or the antenna (Graham and Epstein 1985). In experiments with two stepping legs, either two treadmills or one treadmill and one slippery surface were used. These data were pooled for the analysis.

### Electrophysiology

Extracellular hook electrodes (Schmitz et al. 1988) were used to record the activity of different MN pools. The MN pool innervating the *protractor coxae* muscle (protractor) was recorded from the *nervus lateralis 2* (nl2), the antagonistic MN pool innervating the *retractor coxae* muscle (retractor) was recorded from the *nervus lateralis 5* (nl5) (nomenclature according to Marquardt 1940). In some experiments with freely moving legs, additional electromyographic recordings were performed from flexor tibiae muscles in the front or hind leg according to the established procedures (Schmitz et al. 2015). The measured signals from extracellular recordings were amplified and filtered using an isolated low noise preamplifier (MA101) and a 4-channel amplifier/ signal conditioner (MA102, Electronics Workshop, Animal Physiology, University of Cologne, Germany; gain 1000x, filter: 250 Hz low-cut and 3.5 kHz high-cut). The signals were then digitized by an analog-to-digital converter (Micro 1401 mkII, CED, Cambridge, UK) with a sampling rate of 12,5 kHz and monitored using the Spike2 software (7.20, CED, Cambridge, UK).

### Stimulation of trochanteral/femoral campaniform sensilla

An Aurora Dual-Mode Lever System (Model 300B, Aurora Scientific Inc., Ontario, Canada) was integrated into the aforementioned setup to stimulate tr/fCS during stepping. The system consisted of a lever arm, which applied force through pulling, and a Dual-Mode Lever System, which controlled the applied force output. One end of a strong string was tied to a central position of the trochantero-femur of the middle leg and the other end was attached to the lever arm so that the lever arm could apply force to the leg (Fig. 1A). The Lever System was set on maximum length output, which makes the lever arm fully dependent on the force output of the system. The force applied to the leg and the duration of the stimulation of the tr/fCS were controlled via a control panel on the PC through a custom-written Spike2 script. To stimulate tr/fCS by slight posterior bending of the femur, the lever arm was placed behind the abdomen of the stick insect so that the string was parallel to the body axis. The force output was set to approx. 0.5 mN to keep the string under slight tension while the leg performs stepping movements. Additional force was applied using ramp-and-hold stimuli and the stimulus strength during the holding phase was set to 40 mN. The duration of the holding phase varied between 200 ms and 600 ms, with 400 ms used in most of the experiments. The ramp was set to a velocity of 62.5 mN per 100 ms (Fig. 1B).

### Data Analysis

Stored data was analyzed and plotted using Spike2 (7.20, CED, Cambridge, England) and MATLAB (R2019b, MathWorks, Natick, MA, USA). Bar charts were made with Excel (2016, Microsoft, Redmond, WA, USA) and OriginPro (2021b; OriginLab Corporation, Northampton, MA, USA). All presented figures were arranged and modified using CorelDRAW (X8, Corel Corporation, Ottawa, Canada).

Activities of protractor and retractor MNs throughout the step cycle are presented in the form of step cycle histograms, with a step cycle being defined as the time from the onset of one leg stance to the onset of the next leg stance (Fig. 2Aii, Bii). These are presented in angular degrees with a full cycle set to 360°. Onsets of leg stance were marked manually using the treadmill velocity trace as well as built-in functions in Spike2. For this purpose, a threshold was set slightly above the baseline of the treadmill velocity (no treadmill movement), and all time points at which the treadmill velocity surpassed this threshold were marked as stance onsets. Extracellularly recorded action potentials (APs) of protractor and retractor MNs were marked in a similar procedure: a threshold was set for each recording, which was low enough to mark APs but high enough to exclude noise. The APs of protractor and retractor MNs and the onsets of stance were stored in separate memory channels and exported for further analysis in MATLAB. To allow comparison of the MN activity between forward and backward stepping, all APs were summed in bins of 20 ms along the 360° cycle for each step cycle and normalized to the highest value. Data from all step cycles was then pooled and the average (normalized) number of APs was plotted in the final histogram.

**Figure 2:**
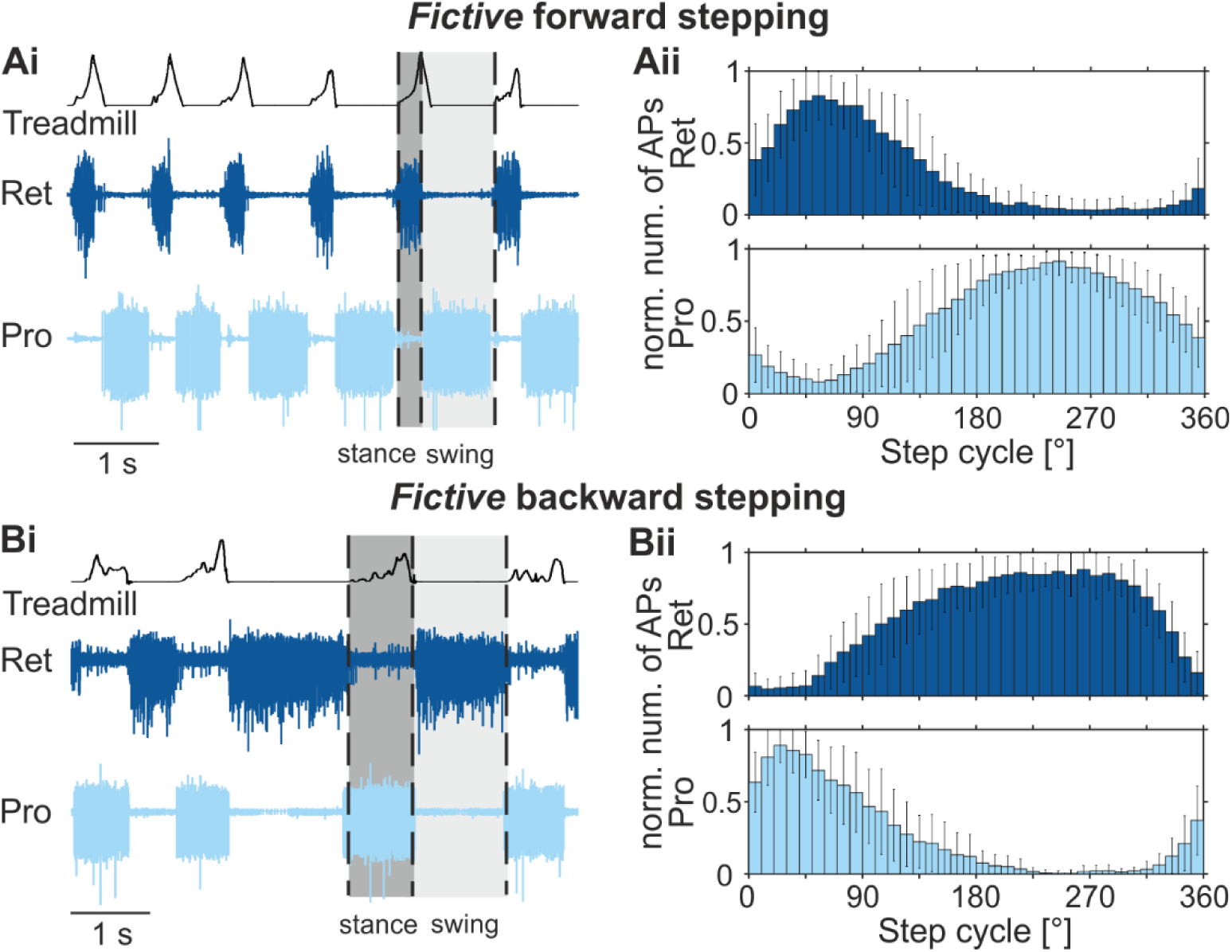
Retractor and protractor motor neuron activity during fictive forward and backward stepping of the middle leg. **Ai, Bi:** Extracellular recordings of retractor (nerve nl5; dark blue) and protractor (nerve nl2; light blue) MNs during fictive forward (**Ai**) and backward (**Bi**) stepping of the middle leg. The upper trace shows the treadmill movement; the ascending slope marks the stance phase, and the decreasing slope and baseline mark the swing phase of the step cycle. To illustrate a step cycle, one stance and swing phase are highlighted in dark grey and light grey, respectively. Transitions from swing to stance and vice versa are indicated for one step cycle by dotted lines. **Aii, Bii:** Histograms showing the normalized number of APs and the standard deviation of retractor (dark blue) and protractor (light blue) MNs during fictive forward (**Aii**) and backward (**Bii**) stepping throughout the step cycle. The step cycle starts with the stance phase and ends with the next stance phase. N=18 animals, n=171 steps in total for fictive forward stepping; N=19, n=185 steps in total for fictive backward stepping. APs: action potentials; MN: motor neuron; Pro: protractor coxae; Ret: retractor coxae.

To analyze the effect of tr/fCS stimulation on protractor and retractor MN activity (Fig. 3Aii, Bii; Fig.4Aii, Bii, Fig. 5Aii, Bii), either the start or the end of the stimulations was marked manually in Spike2. Additionally, all APs of a recording were marked for both protractor and retractor MNs, as described above. These events, i.e., the start or end of the stimulation and the APs, were exported with a 1000 Hz sampling rate and used to plot protractor and retractor activity either around the stimulation start or end in peri-stimulus-time-histograms (PSTH) using a custom-written MATLAB script. For that, APs that occurred in a defined time frame before and after stimulation start or end were summed up in bins of 20 ms width and divided by the number of applied stimuli, resulting in the mean number of APs for each bin.

**Figure 3:**
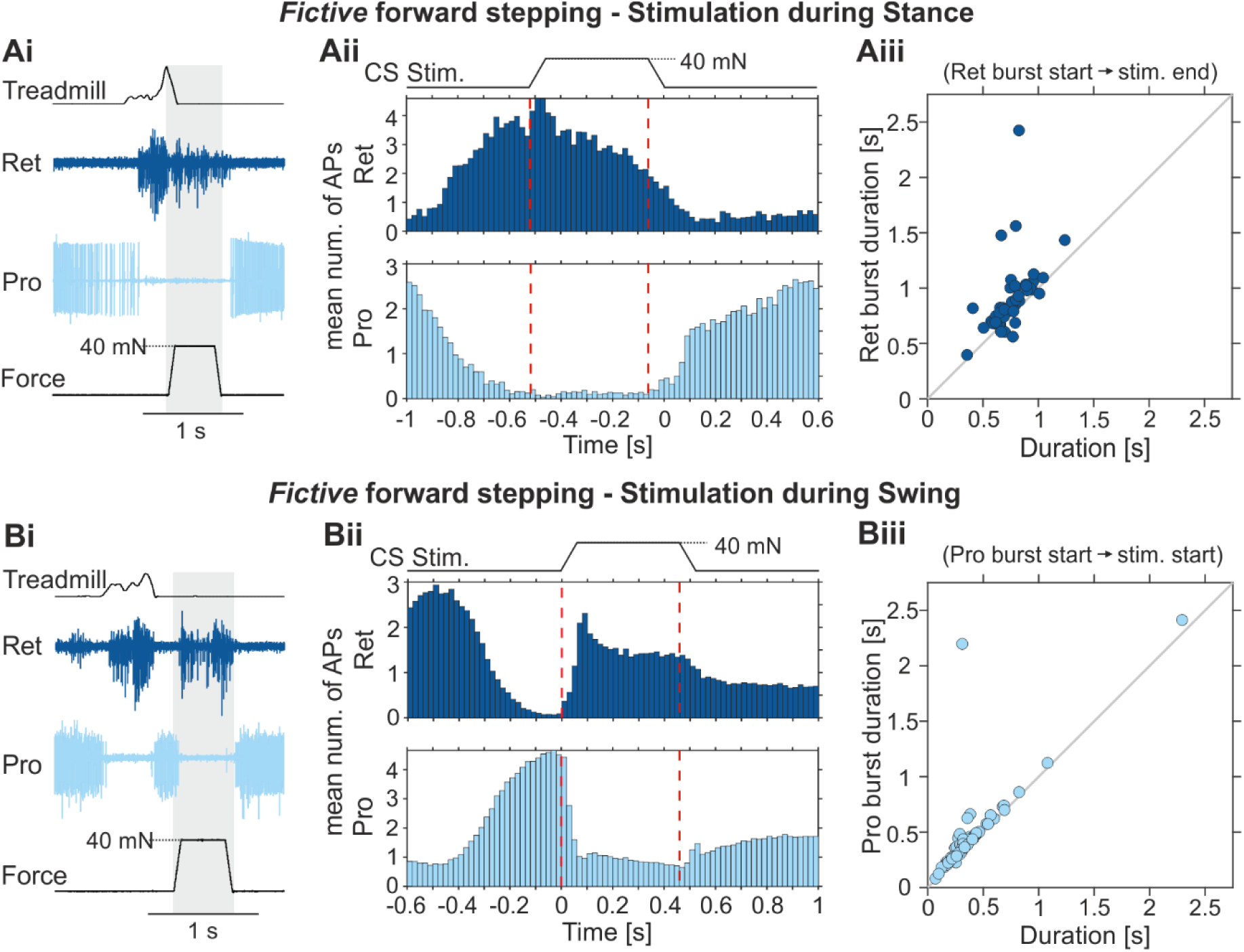
Effects of CS stimulation on protractor and retractor motor neuron activity during fictive forward stepping of the middle leg. **Ai, Bi:** Extracellular recordings of retractor (nerve nl5; dark blue) and protractor (nerve nl2; light blue) MNs during fictive forward stepping of the middle and simultaneous stimulation of load sensors in stance (**Ai**) and swing (**Bi**) phase. The upper trace shows the treadmill movement; the ascending slope marks the stance phase, and the decreasing slope and baseline mark the swing phase of the step cycle. The bottom trace shows the force used to stimulate CS. Load was applied to the leg by pulling it in posterior direction using a ramp-and-hold stimulus with a maximum force of 40 mN and a holding phase of 400 ms. **Aii, Bii**: Histogram showing the mean number of APs of retractor (dark blue) and protractor (light blue) MNs before, during, and after stimulation of CS in stance (**Aii**) and swing (**Bii**) phase. N=8 animals, n=66 stimulations in total during stance, N=19 animals, n=625 stimulations in total during swing. Red dotted lines mark the start of the CS stimulation (loading of the leg) and the end of the stimulation (unloading of the leg). **Aiii**: Dot chart showing the retractor burst duration as a function of the duration of retractor burst start until the end of the stimulation (holding phase), N=5, n=48. **Biii**: Dot chart showing the protractor burst duration as a function of the duration of the protractor burst start until the start of the stimulation. APs: action potentials, N=5, n=68; CS: campaniform sensilla/load sensors; MN: motor neuron; Pro: protractor coxae; Ret: retractor coxae.

To test whether a ramp-and-hold load stimulus applied during the leg’s stance phase extends the burst duration of the stance MN pool for the duration of the stimulation, we plotted the burst duration of the stance MN pool as a function of the time between burst start and end of the load stimulus (Fig. 3Aiii, 5Aiii). For this, we marked the start and end of the stance MN pool burst as well as the end of the stimulation (end of the holding phase) manually in Spike2. These events were stored in memory channels and exported for further analysis with MATLAB.

In addition, to test whether the burst duration of the swing MN pool depends on the load stimulus, we plotted the burst duration of the swing MN pool as a function of time between the start of the burst and the start of the load stimulus (Fig. 3Biii, 5Biii). For this, we marked the start and end of the swing MN pool burst as well as the start of the stimulation (beginning of the ramp) manually in Spike2. These events were again stored in memory channels and exported for further analysis with MATLAB.

To determine the probability of a switch in motor activity during swing phase (Fig. 6), the number of tr/fCS stimulations that induced a switch from swing to stance MN activity was summed up and divided by the total number of stimulations applied during swing for each experimental condition: single middle leg stepping, front and middle leg stepping, hind and middle leg stepping, front, middle and hind leg stepping, and all legs stepping. These data were plotted as bar charts for both forward and backward stepping using OriginPro.

## Results

### Influence of leg CS-stimulation during fictive forward stepping of the single middle leg

The activity of the antagonistic MNs of the ThC joint, the protractor and retractor MNs, was recorded extracellularly in the single leg stepping preparation (Fischer et al. 2001; Rosenbaum et al. 2010). In this preparation, the middle leg can generate quasi-intact stepping movements on a treadmill (Fig. 2) in its vertical plane (Zill et al. 2012) without movement of the coxa in the horizontal plane. During leg stance, the insect moves the treadmill towards its body, reflected by the rising ramp of the treadmill trace in the figures. During leg swing, the treadmill movement stops when the animal lifts and extends its leg over the treadmill to a distal position, where it touches the treadmill again. Then, the next step resumes. In this preparation, motor activity of coxal MNs resembling forward stepping can be systematically elicited by gently touching the abdomen with a paintbrush (Fischer et al. 2001; Rosenbaum et al. 2010), and backward stepping motor activity was initiated by gently touching the antennae or the head with a paintbrush (Graham and Epstein 1985; Rosenbaum et al. 2010). Because the leg is not able to move along the horizontal plane, the motor activities are called “*fictive forward stepping* (*FFS*)” and “*fictive backward stepping (FBS)*” (Rosenbaum et al. 2010).

FFS was characterized by the activity of the retractor MNs during leg stance, while protractor MNs showed little to no activity (Fischer et al. 2001). The opposite was observed during leg swing (Fig. 2Ai). This motor activity during FFS is represented by a step-cycle histogram, which shows the activity in the form of the normalized number of APs of either the protractor or retractor MNs as a function of the step cycle (Fig. 2Aii; N=18 animals, n=171 steps in total). Protractor and retractor MNs showed a rhythmic, alternating motor pattern as described by Fischer et al. (2001) and Rosenbaum et al. (2010). During FBS sequences, on the other hand, protractor MNs were active during leg stance, while retractor MNs stayed mainly inactive. The opposite was observed during leg swing, when retractor MNs showed a high level of activity, while protractor MNs stayed mostly silent (Fig. 2Bi). Again, the motor activity in FBS is illustrated by step cycle histograms for protractor and retractor MN activity (Fig. 2Bii; N=19, n=185). Hence, during FBS, the activity of coxal MN pools was reversed in comparison to forward stepping. These results are in line with the motor patterns described by Rosenbaum et al. (2010, 2015).

Using this setting, we first studied the influence of load signals on the activity of coxal MNs during either stance or swing when the single middle leg exhibited FFS upon tactile stimulation of the abdomen (Fig. 3). First, we tested the load influence during leg stance. Retractor MN activity was increased within the first 20 ms after the start of the load stimulus in the majority of the stimulations, i.e., in ca. 62% of the trials (Fig. 3Ai). This also becomes apparent from the peristimulus-time histograms (PSTH) of coxal motor activity (Fig. 3Aii). Following a slight decay in activity, retractor MNs remained active throughout the holding phase of the stimulus. Retractor activity ceased at the end of the ramp-and-hold stimulus, i.e., upon unloading, while protractor MN activity strongly increased at the same time, reminiscent of a switch in activity between both antagonists (Fig.3Ai, Aii; N=8 animals, n=66 stimulations in total). In ca. 76% of the stimuli applied during leg stance retractor MN activity appeared to be present for the duration of the stimulation (Fig. 3Ai). We tested this by plotting the burst duration of retractor MNs as a function of the time between the retractor burst start and the end of the load stimulus (Fig.3Aiii, N=5; n=48). The result indicates that, in most cases, the duration of retractor activity is determined by load signals starting during leg stance and exceeding leg stance.

The influence of load signals, i.e., CS stimulation, was also tested during leg swing: in almost 67% of the cases, load increase caused a switch from protractor MN to retractor MN activity within 60 ms (Fig. 3Bi). This switch is apparent from the PSTH pooled from all experiments showing the mean number of APs for protractor and retractor MNs aligned to the start of the load stimulus. The activity of both MNs remained relatively unchanged until the tr/fCS stimulation ended, after which the retractor MN activity decreased, while the protractor MN activity increased slightly (Fig. 3Bii, N=19; n=625). We further tested whether the protractor burst duration ends with loading of the leg by plotting the burst duration of protractor MNs as a function of the time between the protractor burst start and the start of the load stimulus (Fig. 3Biii; N=5, n=68). The data indeed show that the protractor burst duration is determined by load signals starting during leg swing and ending with an increase in load. Other responses to load signals were also observed, however much less frequently, i.e., an interruption of protractor MN activity occurring in approx. 7%, and no response in 11% of the cases (data not shown).

The results reported above prompted the conclusion that load stimuli exerted during stance in FFS induce and extend retractor activity for the time of the stimulation. We further tested this by modifying the duration of load stimulation (Fig. 4) using holding times of 200 ms (Fig. 4A; N=2, n=10) and 600 ms (Fig. 4B; N=3, n=13) in addition to the 400 ms that we used before.

**Figure 4:**
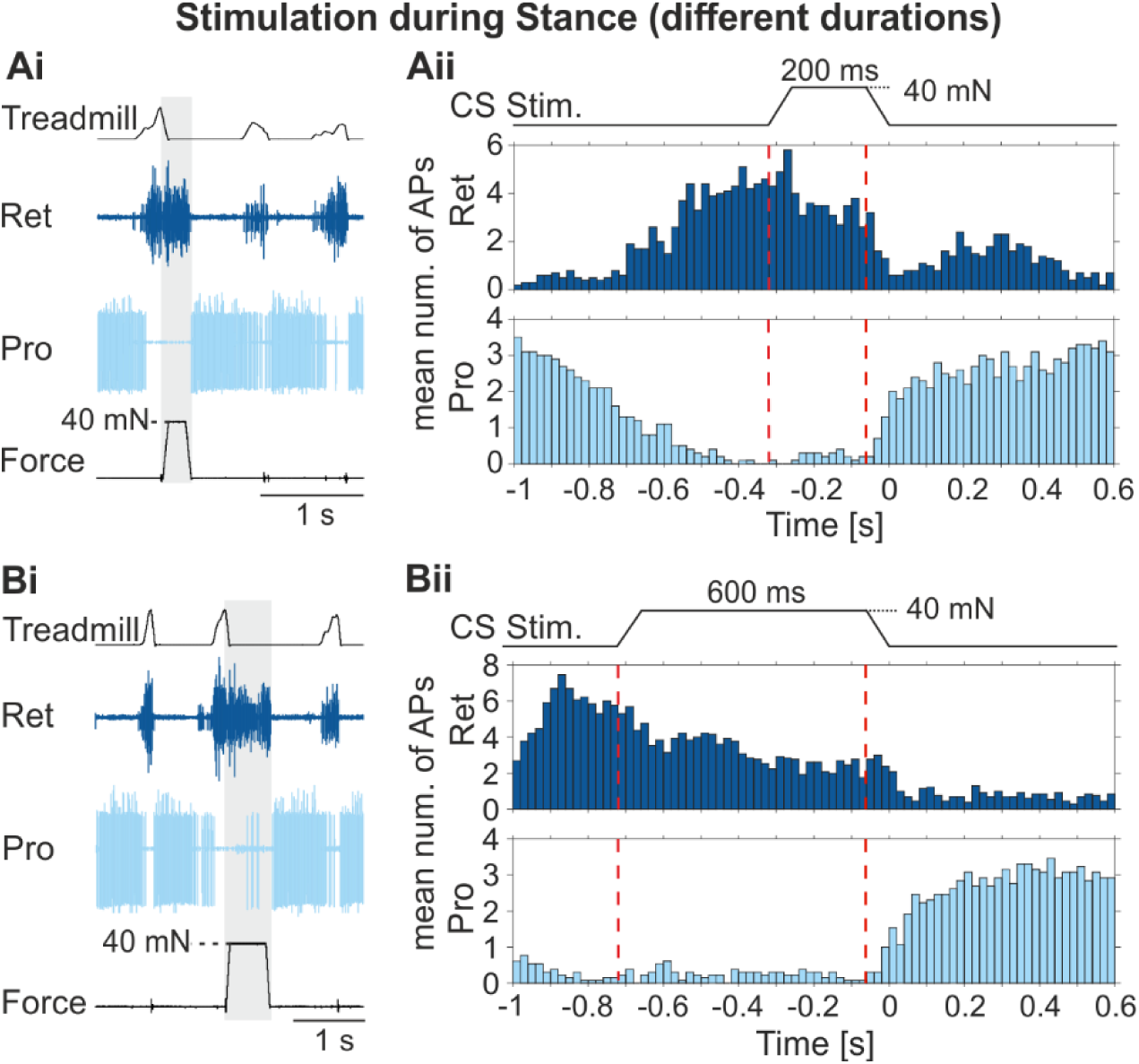
Effects of CS stimulation duration on protractor and retractor motor neuron activity during fictive forward stepping of the middle leg. **Ai, Bi:** Extracellular recordings of retractor (nerve nl5; dark blue) and protractor (nerve nl2; light blue) MNs during fictive forward stepping of the middle and simultaneous stimulation of load sensors in stance with a hold duration of 200 ms (**Ai**) and 600 ms (**Bi**). The upper trace shows the treadmill movement; the ascending slope marks the stance phase, and the decreasing slope and baseline mark the swing phase of the step cycle. The bottom trace shows the force used to stimulate CS. Load was applied to the leg by pulling it in posterior direction using a ramp-and-hold stimulus with a maximum force of 40 mN. **Aii, Bii**: Histogram showing the mean number of APs of retractor (dark blue) and protractor (light blue) MNs before, during, and after stimulation of CS during stance, with a duration of 200 ms (**Aii**) and 600 ms (**Bii**). N=2 animals, n=10 stimulations with a duration of 200 ms, N=3, n=13 stimulations with a duration of 600 ms. Red dotted lines mark the start of the CS stimulation (loading of the leg) and the end of the stimulation (unloading of the leg). APs: action potentials; CS: campaniform sensilla/load sensors; MN: motor neuron; Pro: protractor coxae; Ret: retractor coxae.

As it becomes clear from the PSTHs, retractor MN activity was maintained for the duration of the holding phase, while protractor MNs showed only marginal activity (Fig. 4Aii, Bii). A similar influence was found for the stimulation of load sensors during leg swing (not shown).

### Influence of leg CS-stimulation during fictive backward stepping of the single middle leg

In the next step, we investigated the potential influence of ramp-and-hold stimulation of tr/fCS on the activity of coxal MN pools during FBS, when the activity of protractor and retractor MNs is reversed as compared to FFS (Fig. 2). During leg stance, stimulation of tr/fCS lead to a brief decrease in stance phase MN, i.e., protractor MN activity (Fig. 5Ai; Aii; red arrows). This short decrease in the protractor MN activity was observed in about 33% of the trials. Within about 100 ms, protractor MN activity resumed and was maintained up to the end of tr/fCS stimulation (Fig. 5Aii). A small increase in the activity of retractor MNs was detectable during the decrease in protractor activity, illustrated by the PSTHs (Fig. 5Aii; blue arrow). These PSTHs show the mean activity for the protractor and retractor MNs before, during, and after tr/fCS stimulation, pooled from all experiments and stimulations (N=6, n=33). Interestingly, in almost 70% of the cases, protractor MN activity appeared to be maintained for the time of increased load, exceeding stance duration (Fig. 5Ai, Aii) as shown for retractor MNs in FFS. We further tested this by plotting the burst duration of protractor MNs as a function of the time between the protractor burst start and the end of the load stimulus (Fig. 5Aiii; N=5, n=25). The result indicates that, in most cases, the duration of protractor activity is indeed determined by load signals and not by the duration of leg stance when both happen in parallel. After the stimulation ended, i.e., upon unloading the leg, a switch occurred from protractor MN to retractor MN activity.

**Figure 5:**
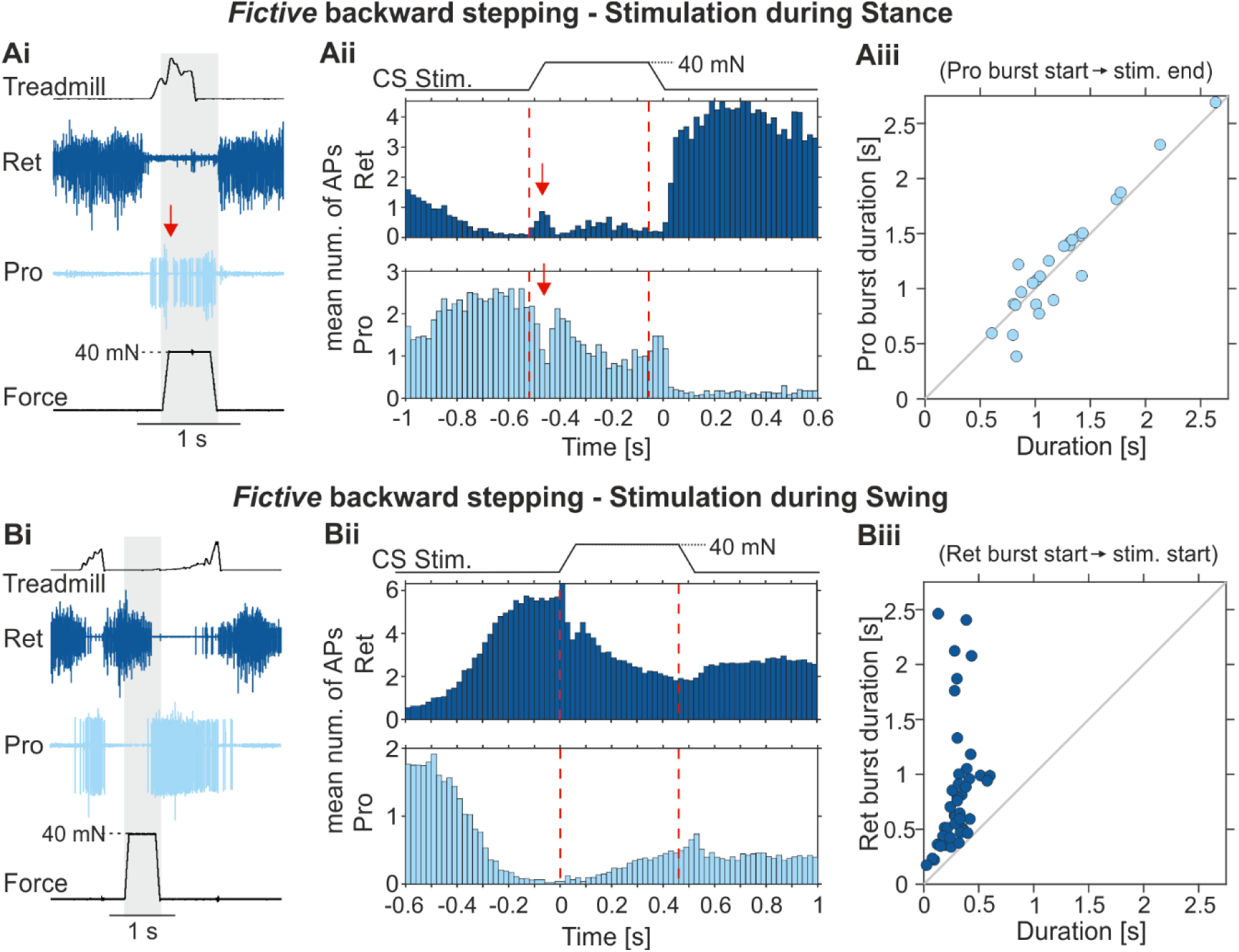
Effects of CS stimulation on protractor and retractor motor neuron activity during fictive backward stepping of the middle leg. **Ai, Bi:** Extracellular recordings of retractor (nerve nl5; dark blue) and protractor (nerve nl2; light blue) MNs during fictive backward stepping of the middle leg and simultaneous stimulation of load sensors in stance (**Ai**) and swing (**Bi**) phase. The upper trace shows the treadmill movement; the ascending slope marks the stance phase, and the decreasing slope and baseline mark the swing phase of the step cycle. The bottom trace shows the force used to stimulate CS. Load was applied to the leg by pulling it in posterior direction using a ramp-and-hold stimulus with a maximum force of 40 mN and a holding phase of 400 ms. **Aii, Bii**: Histogram showing the mean number of APs of retractor (dark blue) and protractor (light blue) MNs before, during, and after stimulation of CS in stance (**Aii**) and swing (**Bii**) phase. N=6 animals, n=33 stimulations in total during stance, N=13, n=205 stimulations in total during swing. Red dotted lines mark the start of the CS stimulation (loading of the leg) and the end of the stimulation (unloading of the leg). Red arrows mark the transient MN responses. **Aiii**: Dot chart showing the protractor burst duration as a function of the duration of protractor burst start until the end of the stimulation (holding phase), N=5, n=25. **Biii**: Dot chart showing the retractor burst duration as a function of the duration of the retractor burst start until the start of the stimulation, N=5, n=47. APs: action potentials; CS: campaniform sensilla/load sensors; MN: motor neuron; Pro: protractor coxae; Ret: retractor coxae.

Interestingly, in FBS and differing from FFS, load signals during leg swing did not markedly affect the activity of protractor or retractor MNs (Fig. 5Bi). Only when quantitatively analyzing MN activity using PSTHs, a slight and brief transient decrease in retractor MN activity upon load increase became apparent (Fig. 5Bii; N=13, n=205). This decrease in activity was observed in ca. 38% of the trials. The activity of the protractor MNs remained mostly unaffected by tr/fCS stimulation (Fig. 5Bii). A switch between the functional swing phase MN pool, i.e., the retractor MNs, and the functional stance phase MN pool, i.e., protractor MNs, was only observed in 6% of the cases in FBS (Fig. 5Bi, Bii). This observation is corroborated by the fact that the burst duration of retractor MNs is more variable and did not depend on the onset of the load stimulation (Fig. 5Biii; N=5, n=47). Furthermore, coxal MN activity appeared not to be markedly affected by an unloading stimulus of tr/fCS during leg swing (Fig. 5Bi, Bii).

### Influence of leg CS-stimulation during fictive forward and backward stepping of the middle leg with different numbers of remaining legs stepping

The results reported above differ from previous reports by Akay et al. (2007). This study reported that load signals from tr/fCS, similar to the ones tested in our study, systematically affect the activity of coxal MNs in opposite ways depending on the walking direction: while in forward walking feedback signals about load increase led to activation of retractor MNs and inactivation of protractor MNs, the opposite was found in backward walking for feedback signals reporting similar load increases. In FBS of the single middle leg, we could not detect a systematic load feedback-induced switch from retractor to protractor activity (cf. our Fig. 5B and Fig. 6). It is this latter finding that we could not verify in the single middle leg preparation. One difference between the experimental approaches of Akay et al. (2007) and ours is that in our case, only the stimulated leg was generating stepping movements, and all other legs had been removed, while in their study, the other five legs were actively performing stepping movements. This might point towards the possibility that intersegmental influences from other stepping legs are necessary to gate load feedback signals in controlling the activity of coxal MNs. To test for that, we compared the probability of a load-induced switch from swing to stance coxal MN activity in both forward and backward walking with different numbers and combinations of participating legs (Fig. 6).

**Figure 6:**
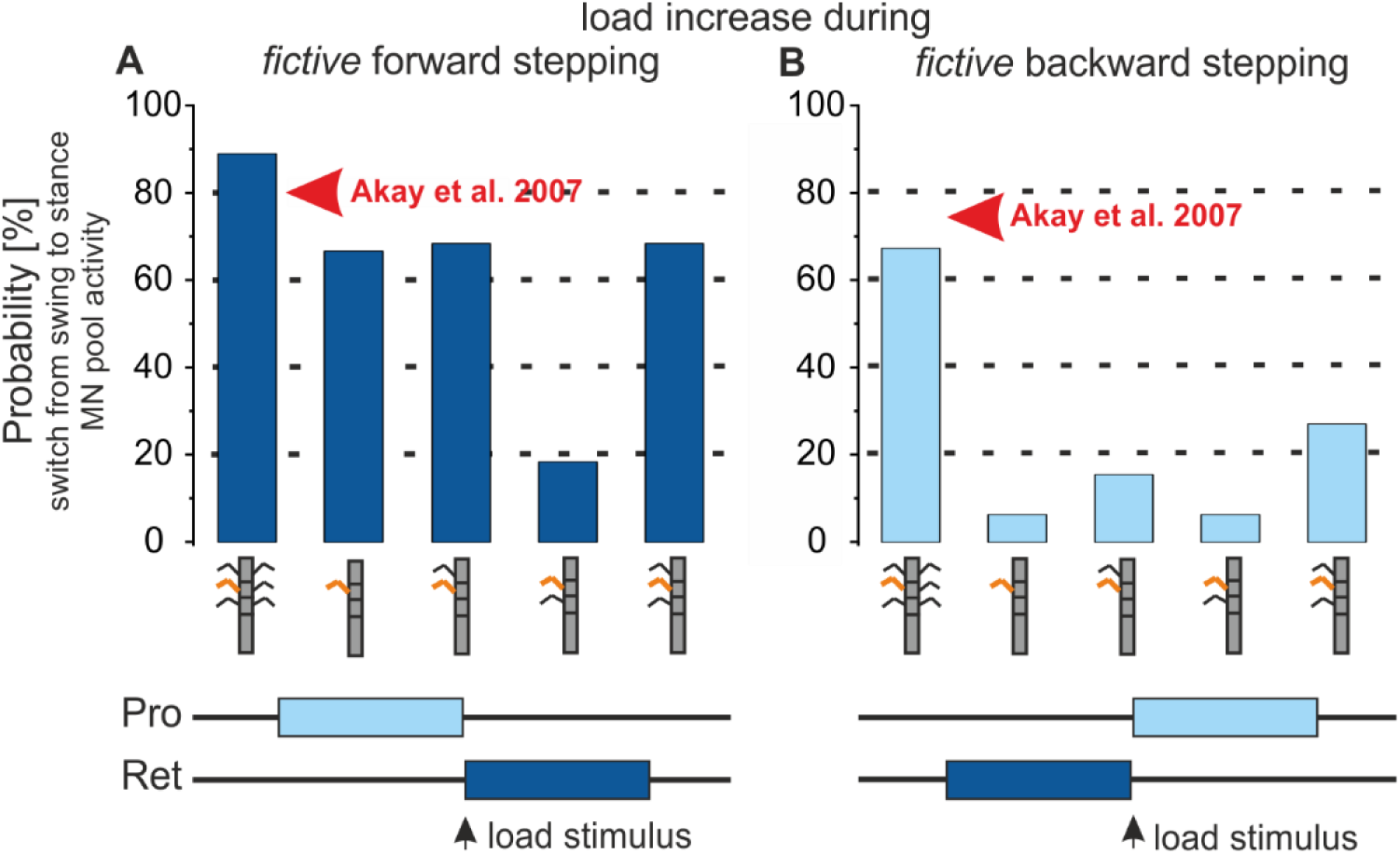
Influence of neighboring legs on load-induced swing-to-stance transition in protractor and retractor motor neurons during fictive forward and backward stepping. Bar plots showing the probability of a switch from swing to stance MN activity in the middle leg induced by stimulation of load sensors during the swing phase in fictive forward (**A**) and backward (**B**) stepping legs. Each bar represents the probability of a switch with different numbers of stepping legs. From left to right: all six legs (forward: N=4 experiments, n=73 stimulations in total; backward: N=3, n=49); only the middle leg (forward: N=19, n=625; backward: N=13, n=205); the middle leg and the ipsilateral front leg (forward: N=4, n=43; backward: N=4, n=45); the middle leg and the ipsilateral hind leg (forward: N=11, n=114; backward: N=11, n=110); the middle leg and the ipsilateral front and hind legs (forward: N=4, n=57; backward: N=5, n=48). Schematic at the bottom illustrates the expected switch from protractor (light blue) to retractor (dark blue) MN activity during fictive forward stepping and from retractor (dark blue) to protractor (light blue) MN activity during fictive backward stepping. The red arrow indicates the probability of a switch when the middle leg stump is stimulated while the 5 remaining legs walk forward or backward on a slippery surface, as reported by Akay et al. 2007. MN: motor neuron; Pro: protractor coxae; Ret: retractor coxae.

In animals with all legs, both in forward and backward stepping conditions, load increase in the stimulated leg resulted in the majority of the cases (89% in forward and 67% in backward walking) in a switch from swing to stance motor activity in coxal MNs, that is a switch from protractor to retractor MNs in forward walking and from retractor to protractor in backward walking. This result parallels the observations reported by Akay et al. (2007; their Fig. 6B). The probability of this switch to occur decreased for both walking directions when the number of legs being active was reduced: for the forward stepping condition, the probability decreased from 89% with five legs stepping to 67% for the single middle leg preparation, and to around 60% when either both front and hind leg or only the front leg was present and stepping. However, when only the ipsilateral hind leg was present and actively stepping, the probability markedly decreased to around 18% (Fig. 6A).

For backward stepping sequences the results differed: in this condition the probability of sensory stimulation reporting load increase to induce a switch from swing to stance motor activity, i.e., from retractor to protractor activity, was strongly reduced in all conditions where no contralateral legs were present and stepping, and only reached about 27% with both the ipsilateral front and hind leg being present and stepping (Fig. 6B). Hence, these results show that intersegmental signals about leg stepping activity play a role in modulating influences of load signals from the middle leg in a task-specific manner, i.e., activation of protractor MNs for generating the stance motor output for backward stepping.

## Discussion

The present study has shown that local sensory feedback signals about load increase and decrease affect the activity of coxal MN pools in a single-stepping middle leg, both when the leg was generating forward stepping-like motor activity as well as backward stepping-like motor activity, called FFS and FBS. Interestingly, however, load-specific influences differed for stance and swing phases of leg stepping and between both motor activities. Our protocol for load feedback stimulation consisted of four phases, i.e., a ramp-wise increase in load, a holding phase of increased load, a ramp-wise reduction in load, and a subsequent phase without load stimulus. We will discuss the observed influences along the sequence of these phases. (i) In FFS, a ramp-wise increase in load during leg stance led to an immediate transient increase in retractor activity (Fig. 3A). (ii) Ongoing load feedback signals resulted in retractor activity being maintained throughout the load stimulus and protractor remaining inactive with this influence exceeding stance phase duration (Fig. 3Aiii). (iii) Upon load decrease retractor activity was terminated and a switch to protractor activity was observed (Fig. 3A). Additional load during leg swing terminated protractor activity and initiated retractor activity (Fig. 3B). Upon the end of a load stimulus, retractor activity decreased and protractor activity increased, however, no regular switch in activity was observed (Fig. 3B).

These influences confirm, for the actively stepping middle leg, previous findings on the influence of CS feedback in forward stepping when load is increased: initially, continuous stimulation of leg CS at the level of the trochanter and femur was found to keep a leg in continuous stance, i.e., to avoid the transition to leg swing after reaching the posterior extreme position during leg stance (Bässler 1977). Subsequently, Akay et al. (2001) showed that it is specifically load feedback from the fCS promoting and enhancing flexor MN activity during leg stance and load feedback from the trCS affecting the CPG of the ThC joint, promoting retractor activity during stance and terminating leg swing activity of protractor MNs in forward walking (Akay et al. 2004). These latter insights were extended by the study of Akay et al. (2007) in a preparation, in which trCS and fCS were selectively stimulated in a middle leg stump and in an animal generating leg stepping movements with all remaining 5 legs intact. In this preparation, the effects of signals about increasing and decreasing load on the CPG of the ThC joint depended on the walking direction: increased load promoted stance phase activity for both stepping directions, with retractor being promoted during forward and protractor activity being promoted during backward stepping.

### The effect of load depends on the stepping direction

In accordance with previous findings, our data suggest that feedback from CS promotes motor activity that generates stance movement. Moreover, our results further specify these previous insights in the following way: load feedback about increasing and increased load from tr/fCS of the single leg during FFS promoted activity of retractor MNs and inhibited activity of protractor MNs during stance phase of the forward stepping cycle (Fig. 3). Qualitatively the same influence was detectable for these stimuli during leg swing, when feedback signals about increasing load terminated protractor activity and initiated retractor activity. Both influences are reminiscent of the situation found for tr/fCS signals at rest (Schmitz 1993; Haberkorn et al. 2019). Hence, during FFS, feedback signals about load increase promote stance activity in ThC joint MN pools during both step phases in a single leg.

During FBS of the single middle leg, load feedback signals had qualitatively different effects depending on the phase of the stimulus. Signals about increased load during stance were found to promote stance, i.e., protractor activity (Fig. 5). However, this influence only became effective after a brief inactivating influence on protractor MN activity and activating influence on retractor MN activity within the first 100 ms after stimulus start, reminiscent to the influence of similar load feedback signals at rest (Haberkorn et al. 2019). Similar to FFS, the stance, i.e., protractor MN activity promoting influence, was maintained for the time of load increase. Furthermore, feedback signals about load decrease promoted subsequently a transition to leg swing MN activity, i.e., retractor MN activity. Load feedback signals during leg swing had, apart from a small decrease in retractor MN activity, no influence (Fig. 5). A switch to the functional stance MN pool was only observed in 6% of the cases. It is important to mention that although loading of the leg rarely initiated stance activity during swing in FBS, the effect seen during FFS was also absent. Similar as in FFS, CS stimulation during FBS had a promoting effect on stance MN pools (protractor) when the stimulation occurred during stance, but only a minor effect when the stimulation occurred during swing.

In summary, these findings suggest that in FBS of the single leg, pathways for load feedback are selectively open and active during the execution of leg stance motor activity and thus cannot contribute to the initiation of leg stance at the end of swing, as was found for forward stepping (Akay et al. 2004, 2007). Only when five other legs were present and stepping, load feedback signals were found to exert a systematic influence when administered during leg swing (Fig. 6), allowing their contribution to the transition of ThC MN activity from swing to stance.

### Transient influences of load feedback signals in FFS and FBS

Both in FFS and FBS, we observed a transient change in retractor and protractor MN activity at the time of load increase. During FFS, retractor activity increased briefly to levels higher than during the remaining holding phase of the load stimulus (Fig. 3), while in FBS, retractor activity showed similarly increase in activity at the time of protractor activity (Fig. 5). This initial transient response of coxal MNs is reminiscent to the responses reported by Schmitz (1993) for applying load to the posterior direction of the femur in a standing and forward walking stick insect preparation using a double treadwheel. Load stimuli similar to the ones used in this present study were found to result in a transient retractor MN activation and transient protractor inactivation. Thus, our study provides evidence that this transient influence is not dependent on the stepping direction, supporting the conclusion that separate neuronal pathways are involved in the transient and the longer-lasting influence of load signals. On one hand, the transient influence is mediated by reflex-like sensorimotor pathways similar to Schmitz (1993) and Haberkorn et al. (2019). On the other hand, the influence of persistent load feedback signals is mediated by them affecting the phase of activity of the ThC-CPG networks, as described by Akay et al. (2004; 2007). Our data provide evidence of two parallel CS pathways, one rather directly through a reflex-like pathway, and the other through influencing a CPG network.

### Intersegmental influences on load feedback signal effects during FF and FB single-leg stepping

Our study has shown that during FBS, the induction of swing (retraction) to stance (protraction) transition requires the presence and stepping of the neighboring five legs (Fig. 6), comparable to the results shown by Akay et al. (2007). Specifically, the stepping activity of the contralateral legs proved to be relevant for the gating pathway. Load feedback signals induced only in 27% of the cases a switch from swing to stance MN activity when only ipsilateral front and hind legs were present and stepping. On the contrary, probability values reached almost 70% with contralateral legs being present (Fig. 6). However, the nature of the input that determines task-specific processing of load feedback remains unknown. Our data suggest that the stepping of the other legs, specifically the contralateral legs, influences the way the CS input is processed.

Previous studies in fruit flies have identified descending neurons in the brain, so-called moonwalker descending neurons (MDNs), which target VNC interneurons in a distributed fashion and are sufficient to induce backward walking in all six legs upon optogenetic activation (Bidaye et al. 2014; Feng et al. 2020). Further studies on cockroaches revealed that descending motor commands from the brain play a role in modulating reflexes (Mu and Ritzmann 2008a, 2008b; Martin et al. 2015). Our results suggest that, in addition to descending inputs from the brain, intersegmental signals from neighboring legs also contribute to the task-specific processing of sensory feedback. Future studies need to identify the nature of input that determines task-dependent sensory processing, for example, whether the phase of stepping activity in the neighboring legs plays a role, as well as the underlying neuronal mechanism. It is known that signals about the stepping of the neighboring legs can generally affect the local motor activity (Ludwar et al. 2005a, 2005b; Stein et al. 2006; Borgmann et al. 2007, 2009, 2011; Grabowska et al. 2022). Moreover, in locusts, mesothoracic intersegmental interneurons were shown to alter the gain of reflexes in the neighboring hind leg (Laurent and Burrows 1989). In light of our results, it is conceivable that such gain changes through intersegmental signals might also be involved in modulating the effect of load feedback during forward and backward stepping and thus could contribute to motor flexibility. Intersegmental signals could, for example, change the gain or weighting of parallel pathways from CS to protractor and retractor MNs by presynaptic inhibition (e.g., Sauer et al. 1997; Gebehart et al. 2022; cf. Akay et al 2007). Nevertheless, our data show that the direction dependent reversal of feedback influence is contingent upon the stepping direction of the neighboring legs.

In addition to FBS, the results on intersegmental influences on load feedback effects in FFS also showed interesting new results. The effect of load on ThC MN pool activity was only mildly affected by the presence of neighboring stepping legs. The probability of a protractor to retractor MN switch exceeded, in general, 50%, reaching up to 90% when all five neighboring legs were present and stepping. However, when only one other ipsilateral leg was present, the identity of that leg appeared to be relevant. In the presence of the ipsilateral front leg, load increase during protractor activity induced in 60% of the cases a transition to retractor activity. This probability dropped to only 18% when only the ipsilateral hind leg was present. This result appears interesting in the context of earlier studies by Bässler and colleagues, which showed that the inherent stepping directions of the leg muscle control systems of front legs and hind legs differ in stick insects when they are decapitated: front legs are reported to step forward, hind legs are reported to step backward (Bässler et al. 1985). The decrease in probability of load increase to induce a switch in activity from protractor to retractor activity in FFS single middle legs, when the hindleg is present suggests that this sensory influence is gated by stepping activity of the ipsilateral front leg and cannot be mediated by a forward stepping hind leg or may even exert an inhibitory influence. Figure 7 summarizes these findings schematically.

**Figure 7:**
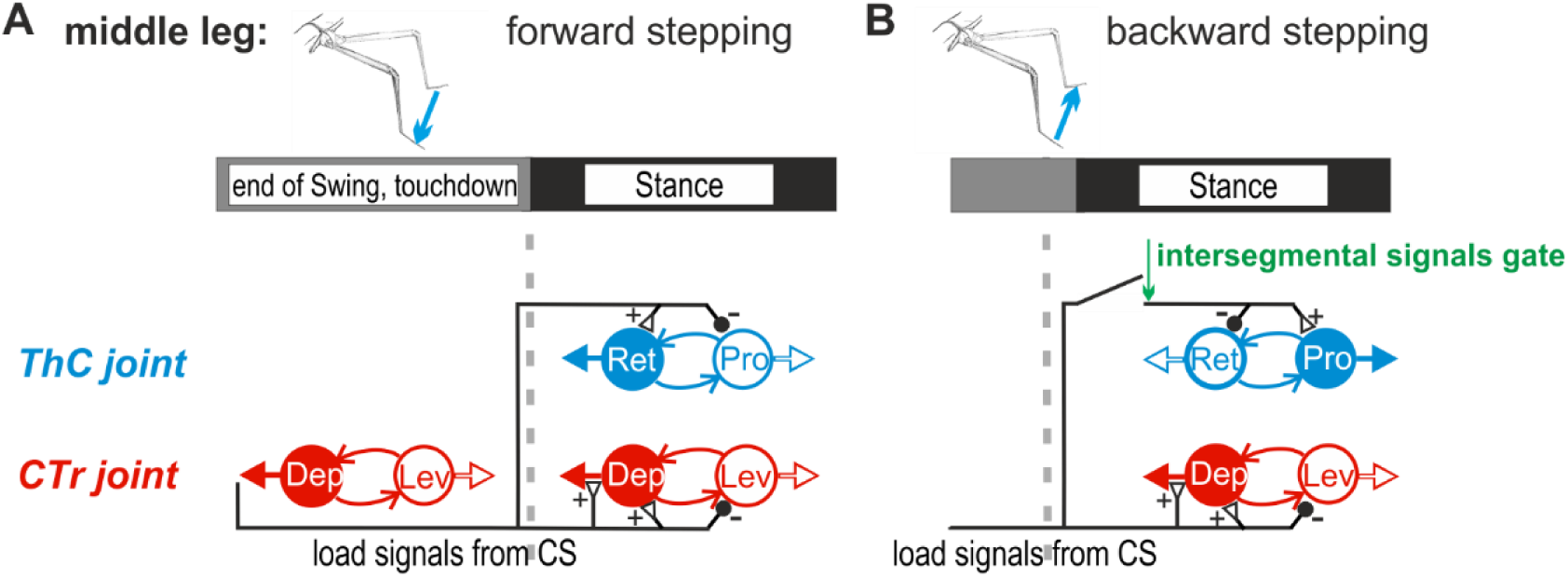
Schematic of the influence of load on the ThC and CTr joint networks during fictive forward and backward stepping (modified after Büschges and Ache, 2025). **A:** Load signals, resulting from depression of the leg at the transition from swing to stance, activate retractor and inhibit protractor activity during fictive forward stepping. They also enhance depressor and inhibit levator activity. **B:** The same load signals reverse their influence during fictive backward stepping as they inhibit retractor and activate protractor activity, presumably through a gating mechanism supported by intersegmental signals. The influence on the CTr joint network does not change. CS: campaniform sensilla; CTr: coxa-trochanter; Dep: depressor trochanteris; Lev: levator trochanteris; MN: motor neuron; Pro: protractor coxae; Ret: retractor coxae; ThC: thorax-coxa.

Taken together, our study shows that local load feedback during either the stance or the swing phases supports the maintenance of or the transition to stance activity regardless of the stepping direction. We further provide evidence that these effects are under the influence of intersegmental neural pathways reporting stepping movements of other legs with a specific role present for the ipsilateral legs in FFS and the contralateral legs in FBS (Fig. 7). Future studies will have to identify the neural mechanisms underlying the task-specific processing of load feedback.

## Acknowledgments

We thank Dr. Till Bockemühl for help with the MATLAB data analysis and Dr. E. Axel Gorostiza for valuable feedback on previous versions of the manuscript. We are grateful to Sima Seyed-Nejadi, Sherylane Seeliger, Michael Dübbert, and Mehrdad Ghanbari for their excellent technical assistance.

## Author Contributions

A.R., A.B., and T.A. conceived and designed the research. A.R., A.Ö., and T.A. performed experiments. A.R. and A.Ö. analyzed data, and A.R. prepared figures. All authors interpreted the results. A.R. and A.B. drafted the manuscript. All authors edited and revised the manuscript and approved the final version.

